# Platelet-Mediated Suppression of T Cell Function Drives Immune Evasion in Triple Negative Breast Cancer through the P-Selectin / P-Selectin glycoprotein ligand-1 Pathway

**DOI:** 10.1101/2025.09.12.675650

**Authors:** Margaret R. Smith-Oliver, Deepa Gautam, Emily M. Clarke, Grace Petrarca, Megan Sullivan, Giovanni Goggi, Milos Spasic, Natalie Kane, Harvey G. Roweth, Ana C Garrido-Castro, Patricia Davenport, Joanna Baginska, Sandra S. McAllister, Elisabeth M. Battinelli

## Abstract

Immune checkpoint inhibitors (ICIs) have demonstrated clinical promise in triple-negative breast cancer (TNBC), yet their effectiveness is often limited by acquired resistance and immune refractoriness. This underscores the urgent need to improve strategies that restore or enhance anti-tumor immunity. Platelets—long recognized for their role in hemostasis—have emerged as key immunomodulators in cancer by interacting with circulating tumor cells, shielding them from sheer stress and immune clearance while actively promoting immune evasion. Here, we uncover a previously unrecognized immunoregulatory pathway whereby platelet-derived P-selectin engages P-selectin glycoprotein ligand-1 (PSGL-1) on T cells, triggering immunosuppressive signaling and promoting T-cell exhaustion. This interaction, identified using in vitro co-culture systems and validated in in vivo mouse models of TNBC, reveals a targetable form of platelet-mediated immune suppression that contributes to ICI resistance. PSGL-1, traditionally known for mediating leukocyte trafficking, functions here as an immune checkpoint receptor, further underscoring the therapeutic relevance of this axis. Together, our findings highlight the P-selectin–PSGL-1 interaction as a novel and targetable mechanism of immune evasion and provide preclinical evidence that its disruption may enhance ICI responsiveness and improve outcomes in TNBC.

**Key Points:**

- Tumor-associated platelets (TAPs) exhaust T-cells through P-selectin/P-selectin glycoprotein ligand-1 binding
- Pharmaceutical blockade of P-selectin using Crizanlizumab, prevents exhaustion and allows T-cell function

## Introduction

Despite significant advances in cancer research and treatment, breast cancer remains one of the leading causes of cancer-related deaths in women [1]. Among its subtypes, triple-negative breast cancer (TNBC) is particularly aggressive, accounting for up to 20% of cases and contributing disproportionately to breast cancer mortality [2, 3]. TNBC lacks expression of estrogen receptor, progesterone receptor, and human epidermal growth factor receptor 2 (HER2), making it refractory to targeted and endocrine therapies and associated with high rates of metastasis and relapse. A growing body of evidence indicates that the tumor microenvironment (TME), a complex network of immune cells, stromal components, extracellular matrix, and inflammatory mediators, plays a vital role in TNBC progression and immune evasion[4–7].

Efforts to activate anti-tumor immunity have led to the development of immune checkpoint inhibitors (ICIs) targeting programmed cell death-1 (PD-1) /programmed cell death ligand 1 (PD-L1) and cytotoxic T-lymphocyte associated protein 4 (CTLA-4), which have revolutionized treatment for many cancers. Of the different types of breast cancer, TNBC—the most aggressive subtype with higher immunogenicity—is uniquely eligible for first-line immunotherapy [8–11]. However, the success of ICIs in TNBC has been modest, with many cases marked by primary or acquired treatment failure [12–15]. While several mechanisms underlie ICI resistance, increasing evidence points to a novel player in immune modulation within the TME, platelets, which extend far beyond their classical role in hemostasis [2, 16]. In particular, tumor-associated platelets (TAPs) engage in direct interactions with immune cells via adhesion molecules such as P-selectin and its ligand P-selectin glycoprotein ligand-1 (PSGL-1), influencing leukocyte trafficking, T cell differentiation, and functional activation [17–19]. Studies have also shown that elevated platelet counts predict poor prognosis in cancer patients [2, 20, 21]. These findings suggest that platelets are not passive bystanders, but active regulators of immune suppression.

Upon interaction with tumor cells, platelets undergo phenotypic reprogramming to become TAPs, adopting a pro-tumoral profile characterized by increased secretion of immunosuppressive cytokines and surface molecule expression [22–25]. TAPs release transforming growth factor-beta (TGF-β), which induces epithelial-mesenchymal transition (EMT) and suppresses effector T cell function [26, 27]. In parallel, platelet-derived PD-L1 is increasingly recognized in both tumor and peripheral compartments as a contributor to T cell inhibition [28]. We previously showed that TAPs form aggregates with tumor cells, shielding them from immune detection by upregulating PD-L1 and phosphorylated epidermal growth factor receptor (pEGFR), thus promoting immune evasion [24]. These immunomodulatory features place TAPs at the intersection of tumor survival and immune escape.

P-selectin is a transmembrane glycoprotein, mainly derived from platelets and endothelial cells, that upon platelet activation translocates from the membrane of α-granules to the cell surface during exocytosis[29, 30]. Under physiological conditions, it facilitates leukocyte adhesion to activated endothelium and platelets through binding to its primary ligand, PSGL-1 [31]. However, in the tumor microenvironment, P-selectin contributes to cancer progression by promoting tumor cell–platelet aggregation, shielding tumor cells from immune surveillance, and facilitating metastatic spread [32–36]. Analysis of The Cancer Genome Atlas (TCGA) has revealed that elevated expression of the gene encoding P-selectin (SELP) is associated with worse overall survival in multiple cancer types, including glioblastoma, melanoma, breast cancer, and colon cancer [37, 38]. These findings highlight the potential immunosuppressive role of P-selectin in cancer and raise the question of whether targeting the P-selectin/PSGL-1 axis could enhance anti-tumor immune responses, particularly in immunologically cold tumors such as TNBC.

Effector CD4+ and CD8+ T cells are crucial mediators of anti-tumor immunity and central targets of immunotherapy. Yet, in the TME, their activity is blunted by suppressive networks involving tumor cells, regulatory T cells (Tregs), and cancer-associated fibroblasts [39]. TAPs may serve as an additional brake on T cell function through their expression of PD-L1, secretion of TGF-β, and direct cell-cell interactions. In particular, the P-selectin/PSGL-1 axis has been implicated in skewing CD4+ T cell differentiation toward Tregs and impairing effector T cell activation in inflammatory disease [40–43]. However, emerging evidence indicates that the P-selectin/PSGL-1 axis plays a role in modulating the immune response in various diseases such as cancer. It has been shown that PSGL-1 acts as a negative regulator of T-cell immune response, therefore facilitating tumor growth in cancerous phenotypes [44]. These platelet-mediated mechanisms likely contribute to T cell exhaustion and diminished ICI efficacy in TNBC.

We hypothesized that TAPs promote immune evasion in TNBC by suppressing CD4+ and CD8+ T cell function through PD-L1 expression and P-selectin/PSGL-1-mediated adhesion, collectively driving T cell exhaustion and immunotherapy resistance. To test this, we utilized in vitro co-culture systems with mouse and human T cells and TAPs, in vivo syngeneic TNBC models, and immunophenotyping to assess T cell cytotoxicity, exhaustion markers, and checkpoint expression. Because P-selectin has been implicated as a driver of TAP–immune interactions, we examined whether its blockade could restore T cell function. To this end, we employed Crizanlizumab, an FDA-approved monoclonal antibody against P-selectin that is currently used to prevent vaso-occlusive crises in patients with sickle cell disease[45, 46]. Together, these approaches enabled us to evaluate the therapeutic potential of P-selectin inhibition with Crizanlizumab as a strategy to overcome TAP-mediated immune suppression in TNBC.

## Methods

### Sex as a Biological Variable and Study Approval

Our study exclusively examined female patients and mice, as approximately 99% of breast cancer cases are in females [47]. Mouse experiments were approved under BWH IACUC protocol #2019N000011; human samples were collected with informed consent under Dana-Farber/Harvard Cancer Center IRB approval and in accordance with the Helsinki Declaration.

### Mice

Female C57BL/6 (Jackson #000664) and B6;129S2-Selp^tm1Hyn/J (Jackson #008437) mice (9 wk) were injected with 2.5 × 10⁵ AT-3 cells (PBS) into the mammary fat pad. Once tumors reached 100 mm³, mice received PBS, anti–PD-1 (150 μg i.p. q4d), RB40.34 (30 μg i.p. 3×/wk), or combination therapy. Tumor volumes were measured by calipers [½(length × width²)] thrice weekly pre-treatment and daily post-treatment; n = 5–10/group [48].

### Platelet Isolation

Mouse blood was collected by cardiac puncture into 3.8% trisodium citrate, diluted 1:1 with Tyrode buffer, and centrifuged (200 × g, 10 min, RT) for PRP. Prostaglandin E1 (1 μM; Sigma #P5515) was added, PRP spun (400 × g, 10 min), and pellets resuspended in serum-free RPMI or TCM at 2 × 10⁸/mL (used <3 h). Human blood (sex-matched healthy donors or metastatic TNBC patients not on platelet inhibitors) was processed similarly: PRP (200 × g, 10 min, RT), PGE1 (Sigma #P5515-1MG, 1:50), centrifuged (400 × g, 10 min), washed in platelet buffer (20 mM HEPES, 138 mM NaCl, 2.9 mM KCl, 1 mM MgCl₂, 0.36 mM NaH₂PO₄, 100 mM EGTA, 5 mM glucose, pH 7.4), and resuspended at 2 × 10⁸/mL.

### Flow cytometry

Platelets (2 × 10⁸/mL) were stained under three conditions: resting, TAPs, or CRP-activated (Pplus Medical #CRP-A0.5-WIN03, 1 μg/mL). Antibodies included CD62P (RB40.34), CD274 (10F.9G2), Galectin-9 (RG9-35), I-Ab (AF6-120.1), CD86 (GL1), CD155 (TX56), Galectin-3 (M3/38). Samples were fixed in 2% paraformaldehyde (Thermo #J19943-K2) and analyzed on Cytek with FlowJo v10.

Mouse spleens were processed into single-cell suspensions (RPMI + 10% FBS, 70 μm strainer; Thermo), RBC-lysed, and stained with Zombie NIR (BioLegend #423105) plus antibodies (**Supplementary Tables 1–2**) [49]. Human PBMCs (10 mL blood, Ficoll separation, RBC lysis) were stained similarly (Supplementary Table 3). Intracellular staining used fixation/permeabilization kits (Thermo Fisher #00-5523-00 or BioLegend) with markers including IFN-γ, Grz-B, TOX, TCF-1, FOXP3. All acquired on Cytek/FlowJo; gating in **Supplementary Figures 1–2**.

### T cell isolation and activation

Mouse CD3⁺ cells were isolated (EasySep #19851) from AT-3–bearing spleens, cultured in RPMI + 10% FBS, 10 mM HEPES, 100 U/mL pen/strep, and activated with IL-2 (Thermo #212-12-20UG) and Dynabeads CD3/CD28 (Thermo #11456D, 1:1) for 48 h. Human CD3⁺ cells were enriched from PBMCs (EasySep #17951), cultured similarly, and activated with IL-2 (PeproTech #200-02) and Dynabeads CD3/CD28 (Thermo #11131D).

### T Cell–Platelet Co-culture and Cytotoxicity

CD3⁺ T cells were cultured for 48 h in serum-free RPMI with: (a) T cells alone, (b) TCM, (c) resting platelets, or (d) TAPs (T cell:platelet ratio 1:250 [42]. For cytotoxicity, T cells were co-cultured with AT-3 cells (3,000/well, seeded 24 h prior) at 10:1 for 16 h. Apoptosis was measured by caspase-3/7 (Thermo #C10423), Zombie NIR, and CD3.

### Cell Culture and Tumor cell conditioned media (TCM) preparation

AT-3 (Sigma SCC178; RRID:CVL_VR89), E0771 (ATCC CRL-3461), and MDA-MB-231 (ATCC HTB-26) cells were maintained in DMEM + 10% FBS + 1% pen/strep. For TCM, confluent AT-3 or MDA-MB-231 cells were washed, incubated 24 h in serum-free RPMI, centrifuged (1000 × g, 5 min), and supernatants stored at −80 °C.

### Statistical analysis

GraphPad Prism v10 was used. Unless otherwise noted, Student’s *t*-test was applied. Significance as *, P< 0.05; **, P< 0.01; *** P< 0.001.

## Results

### Tumor associated platelets (TAPs) Impair Effector T Cell Function and Promote Exhaustion

To determine whether tumor-activated platelets (TAPs) contribute directly to immune evasion, we developed a platelet–T cell co-culture system. For each experiment, platelets were isolated from two healthy mice; platelets from one mouse were kept resting in serum-free RPMI, while platelets from the other were cultured for 10 minutes in AT-3 tumor cell–conditioned medium (TCM) derived from the murine TNBC cell line AT-3 to generate tumor-activated platelets (TAPs). Building on our previous work demonstrating that platelets become activated in TCM media to become TAPs, we used this system as an in vitro model to study TAP–T cell interactions [25, 50]. T cells were isolated from AT-3 tumor–bearing mice two days prior to the experiment, activated with CD3/CD28 and IL-2 for 48 hours, and then co-cultured in serum-free RPMI for 48 hours under one of four conditions: (i) T cells alone, (ii) TCM alone + T cells, (iii) resting platelets + T cells, or (iv) TAPs + T cells (**Figure 1A**). Flow cytometric analysis using a 20-antibody panel revealed that co-culture with TAP resulted in a significant decrease in expression associated with effector potential markers on both CD4^+^ and CD8^+^ effector T cells compared T cells that had been co-cultured with resting platelets or TCM alone (**Figure 1B&1C**). Importantly, neither TCM alone nor resting platelets (i.e., not exposed to TCM) reduced T cell functional markers, indicating that platelet exposure to TAPs is required to elicit these changes.

**Figure 1.**
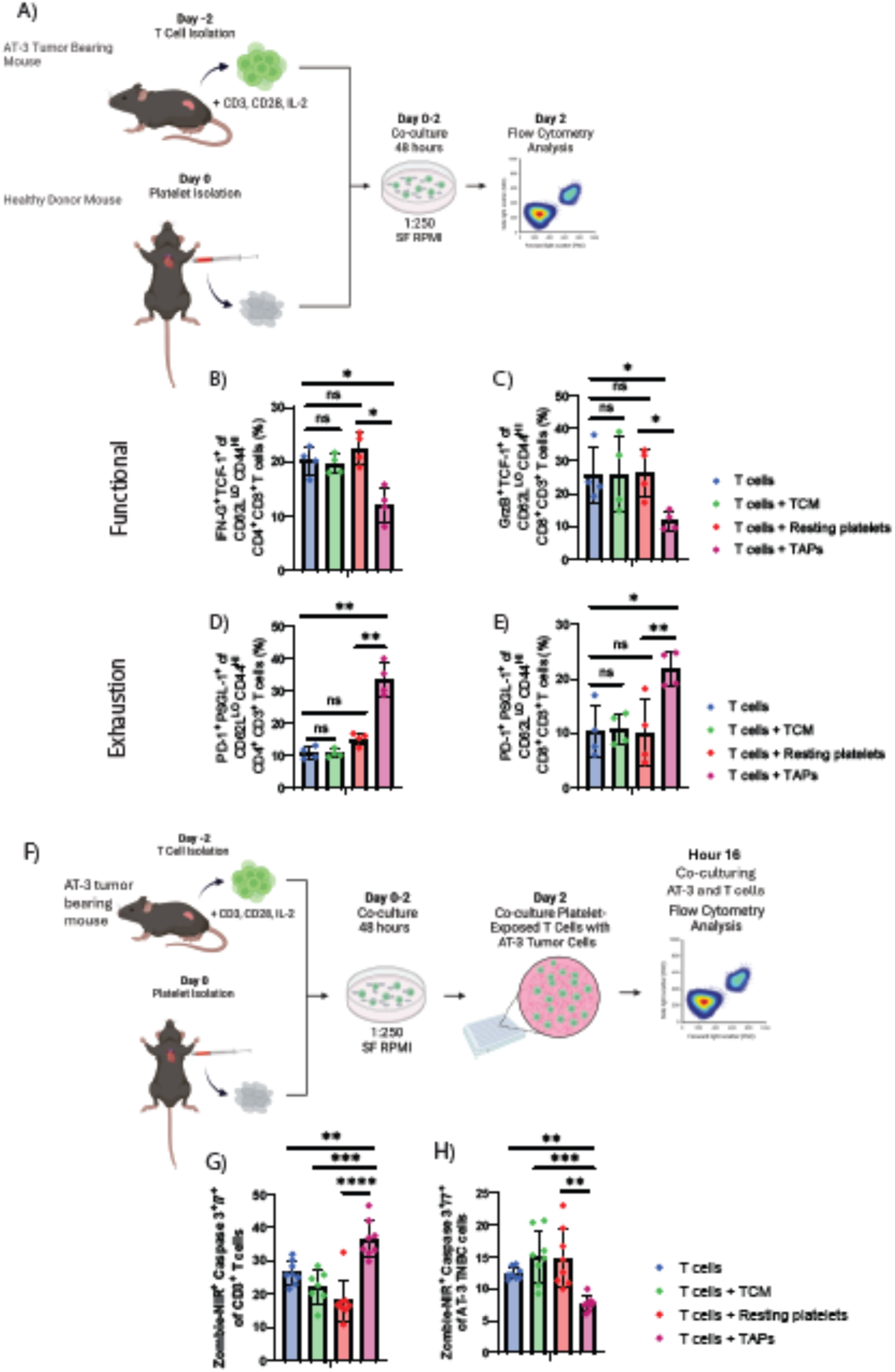
TCM-conditioned platelets exhaust CD4+and CD8+Effector T cells. A) Schematic of isolation and T cells co-cultured with platelets experiment. Flow cytometry used to quantify functionality of (B) CD4+and (C) CD8+effector T cells and exhaustion of (D) CD4+and (E) CD8+effector T cells. (F) Schematic of T cell and AT-3 co-culture experiment. Levels of Zombie-NIR caspase 3/7 dye detected over 48 hours. Flow cytometry to detect death in G) CD3+T cells and H) AT-3 tumor cells. *, *P* < 0.05; **, *P* < 0.01; ***, *P* < 0.001.

To identify potential mediators of platelet-induced T cell suppression, we examined platelet surface markers previously implicated in activation, adhesion, and immune modulation. These included checkpoint ligands (PD-L1, CD155), adhesion molecules (P-selectin, CD86), galectins (Galectin-3, Galectin-9), and the antigen presentation molecule I-Ab. The AT-3-TAPs -activated platelets exhibited elevated expressions of PD-L1, P-Selectin, Galectin-3, Galectin-9 and I-Ab compared to resting or collagen-related peptide (CRP)-activated controls (**Supplementary Figure 2A-G**), suggesting multiple potential inhibitory interactions with T cells [51–56]. Given that P-Selectin is a principal mediator of platelet adhesion and PD-L1 is a well-established immune checkpoint, and based on our lab’s previous investigations of these molecules, we prioritized them to interrogate platelet-mediated T cell suppression while limiting the scope to the most functionally relevant targets [20, 24, 36, 57–59]. We then looked for their cognate receptors on T cells. T cells co-cultured with TAPs showed markedly increased expression of the immune checkpoint and exhaustion markers, Programmed cell death protein 1 (PD-1) and P-selectin Glycoprotein Ligand 1 (PSGL-1), on both CD4⁺ and CD8⁺ effector T cells compared to T cells that were cocultured with TCM alone, resting platelets, or alone without platelets (**Figure 1D&1E**).

To assess cytotoxic capacity, T cells previously co-cultured for 48 hours were incubated with AT-3 tumor cells in the presence of a caspase 3/7 death dye (**Figure 1F**). T cells exposed to TAPs had significantly increased apoptosis and, conversely, were less effective at inducing AT-3 tumor cell death compared to controls (**Figure 1G–H**). These data indicate that TAPs directly impair T cell function and promote exhaustion. Taken together, these findings suggest that TAP-mediated T cell suppression may facilitate tumor immune escape

### Human T Cells Recapitulate Platelet-Mediated Dysfunction

To evaluate the translational relevance of our murine findings, we repeated the co-culture experiments using human T cells from healthy donors. After activation, T cells were co-cultured with MDA-MB-231-TCM alone, resting human platelets, or MDA-MB-231 TAPs (**Figure 2A**). Similar to the murine data, TAPs significantly decreased expression of functional markers on CD4⁺ and CD8+ effector T cells while resting platelets or TCM alone had no effect (**Figure 2B– C**). Exhaustion markers PD-1 and PSGL-1 were significantly upregulated on CD4⁺ and CD8⁺ effector T cells in the presence of TAPs (**Figure 2D–E**). These results support that TAP can suppress T cell function and induce exhaustion in both mouse and human systems, suggesting their immunosuppressive role in a tumor setting.

**Figure 2.**
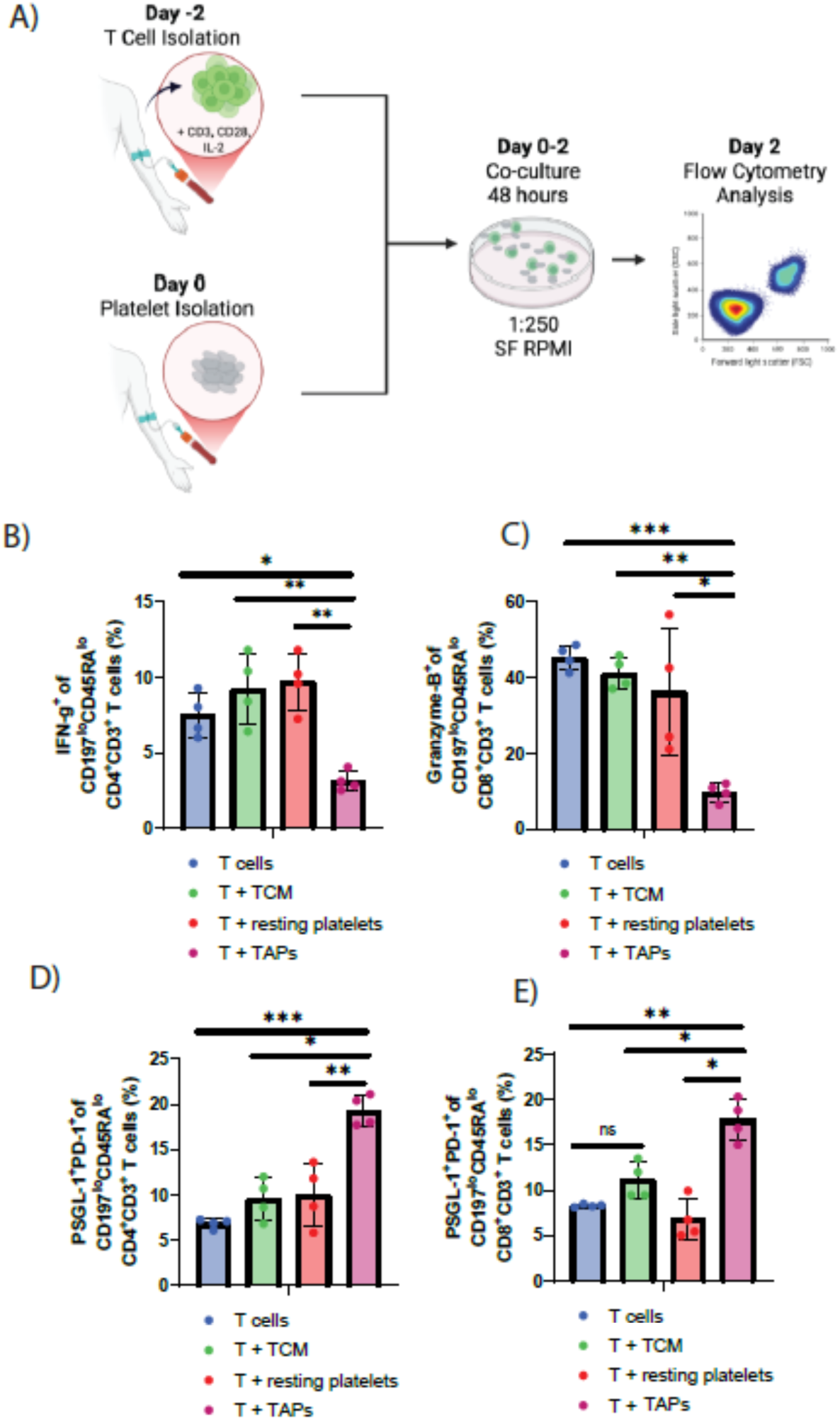
TCM-conditioned platelets exhaust CD4+and CD8+Effector T cells in human. A) Schematic of isolation and T cells co-cultured with platelets experiment. Flow cytometry used to quantify functionality of B) CD4+ and C) CD8+ effector T cells and exhaustion of D) CD4+and E) CD8+effector T cells in TNBC co-culture with Crizanlizumab. *, *P* < 0.05; **, *P* < 0.01; ***, *P* < 0.001.

### Loss of P-Selectin Partially Restores T Cell Functionality and Reduces Exhaustion

Since platelets interact with T cells through the P-Selectin/PSGL-1 axis[60], we hypothesized that P-Selectin is a key mediator of T cell dysfunction. Using platelets from P-Selectin knockout mice (B6;129S2-*Selp^tm1Hyn^*/J), we found that TAPs KO-P-Selectin did not express surface P-Selectin yet displayed elevated PD-L1 levels compared to those from wildtype C57BL/6 mice (**Figure 3A-B**). T cells co-cultured with TAPs KO-P-Selectin exhibited partially restored expression of effector-associated markers in both CD4⁺ and CD8⁺ subsets compared to TAPs.

**Figure 3.**
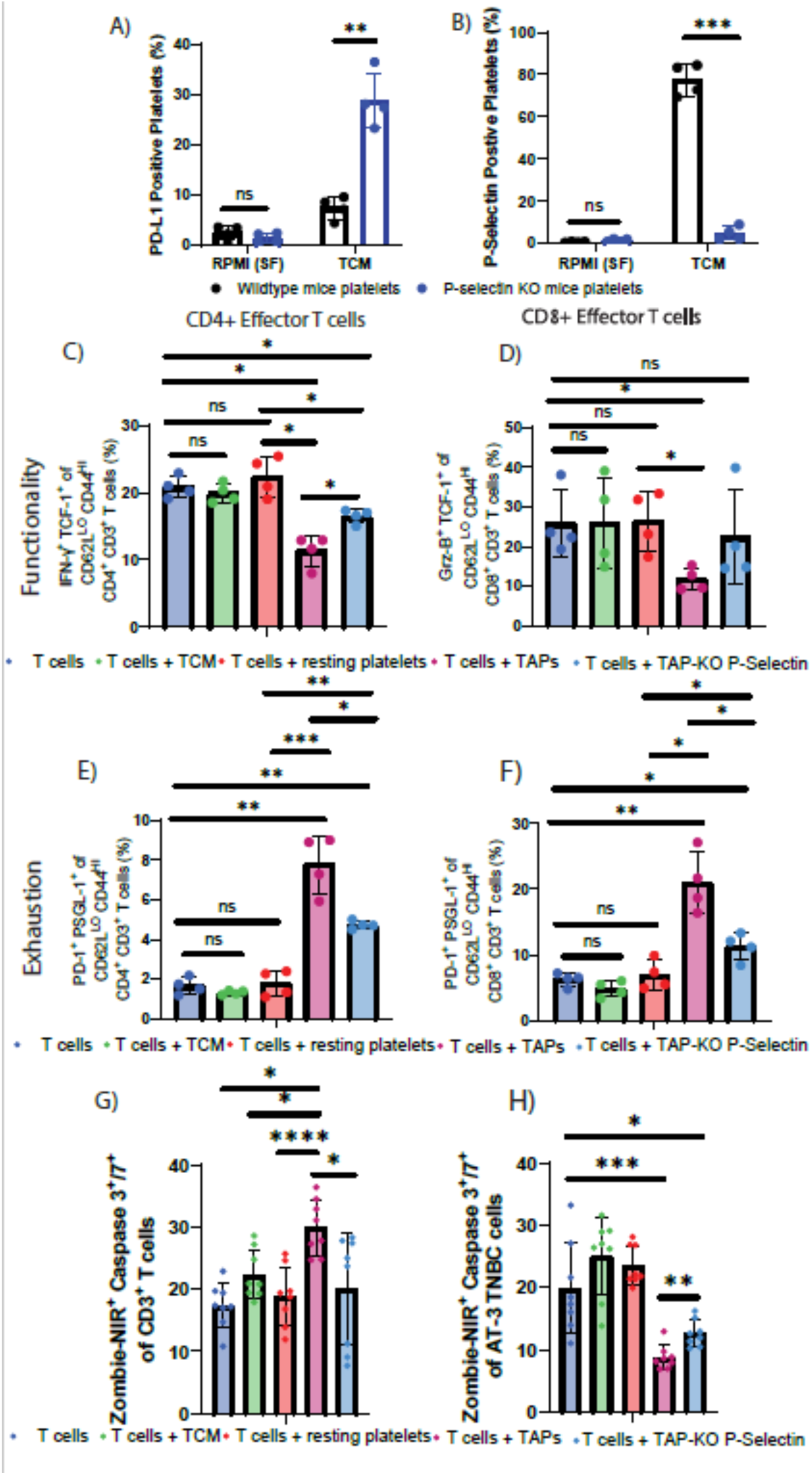
TCM-activated KO platelets from P-Selectin knockout mitigates T cell dysfunction. Flow cytometry used to quantify TCM-activated KO platelet levels of A) PD-L1 and B)P-Selectin, functionality of T cells co-cultured with TCM-activated P-Selectin KO platelets in both C) CD4⁺ and D) CD8⁺ effector T cells and exhaustion of E) CD4+ and F) CD8+ effector T cells. Levels of Zombie-NIR caspase 3/7 dye detected over 48 hours. Flow cytometry to detect death in G) CD3^+^ T cells and H) AT-3 TNBC cells co-cultured with TCM-activated P-Selectin KO platelets. *, *P* < 0.05; **, *P* < 0.01; ***, *P* < 0.001.

(**Figure 3C–D**). Correspondingly, T cell exhaustion markers PD-1 and PSGL-1 were reduced in KO co-cultures compared to T cells co-cultured with TAPs (**Figure 3E–F**), indicating that the loss of P-Selectin mitigates T cell dysfunction. T cell cytotoxicity assays confirmed increased AT-3 tumor cell killing, and reduced T cell death, when using T cells previously co-cultured with TAPs KO-P-Selectin versus TAPs (**Figure 3G–H**). These results highlight a critical role for P-Selectin in platelet-mediated T cell suppression, with its absence restoring partial immune competence.

### Crizanlizumab Blocks Platelet–T Cell Interaction and Preserves T cell Functionality

Given the central role of P-Selectin in platelet-mediated immune modulation and evidence that P-Selectin KO mice exhibit reduced TNBC tumor growth and decreased regulatory T cell infiltration [61], we next tested whether its pharmacological inhibition could reverse T cell suppression with Crizanlizumab, an FDA-approved anti–P-Selectin antibody [62], significantly reduced the formation of platelet–T cell aggregates in both CD4⁺ and CD8⁺ T cells at 0.8 mg/mL, a key interaction through which platelets suppress T cell effector function (**Figure 4A-B**) [63, 64]. Using the RB40.34 antibody, the murine equivalent to Crizanlizumab, we co-cultured T cells with TAPs in the presence or absence of inhibitor. Both Crizanlizumab and RB40.34 preserved T cell functionality as assessed by preservation in effector-associated marker (**Figure 4C–D**) and prevented exhaustion marker upregulation (**Figure 4E–F**). Furthermore, inhibition of P-Selectin during platelet co-culture restored some of the tumor cell–killing capacity of T cells, as measured in cytotoxicity assays (**Figure 4G–H**). These findings demonstrate that pharmacological blockade of P-Selectin can effectively protect T cells from platelet-induced suppression.

**Figure 4.**
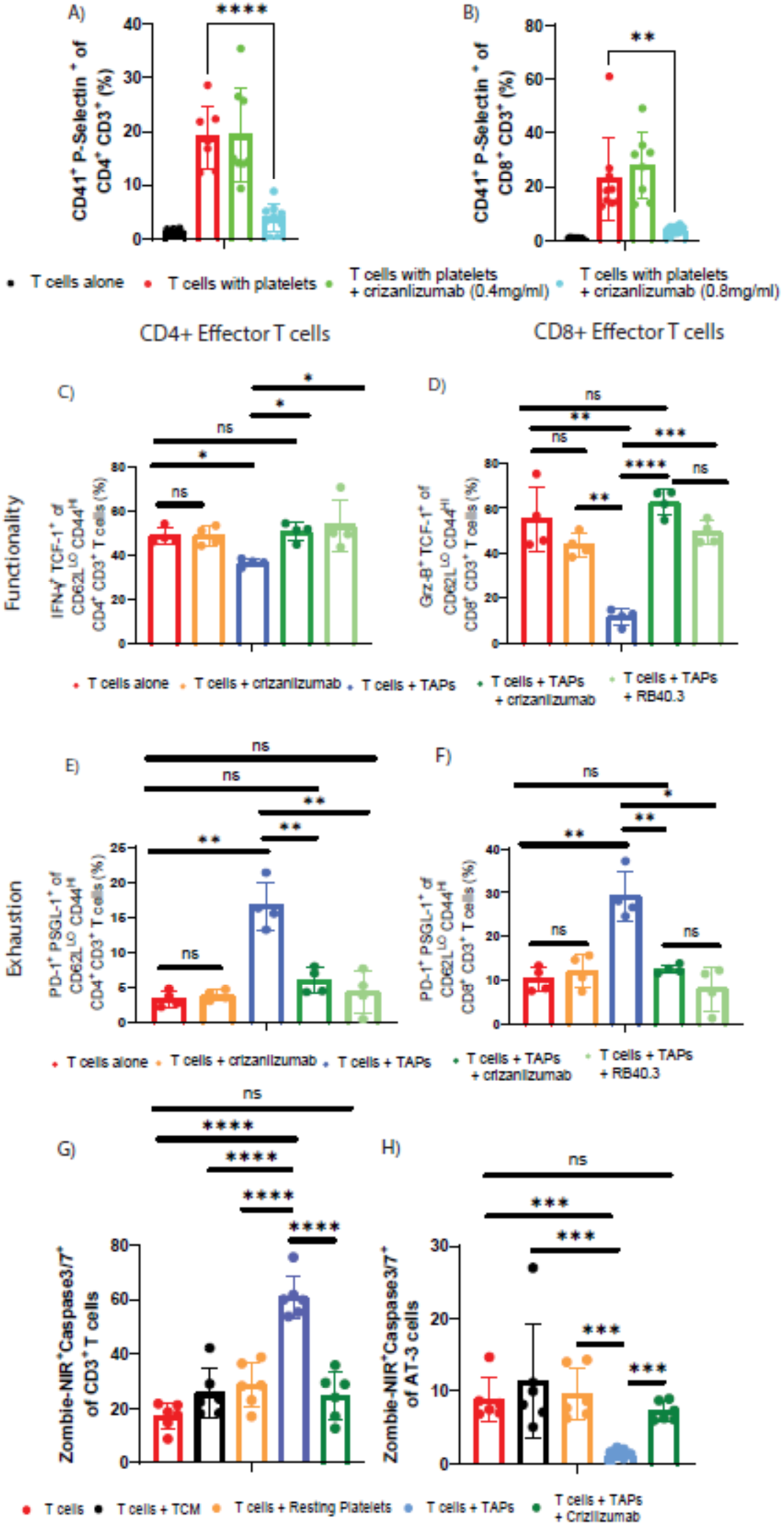
T cell function preserved with blocked platelet-T cell interaction through introduction of Crizanlizumab. Flow cytometry quantification of platelet–T cell binding in both A) CD4⁺ and B) CD8⁺ effector t cells population. Quantified functionality of T cells co-cultured with TCM-activated platelets and Crizanlizumab and RB40.3 in both C) CD4⁺ and CD8⁺ D) effector T cells and exhaustion of E) CD4+ and F) CD8+ effector T cells. Levels of Zombie-NIR caspase 3/7 dye detected over 48 hours. Flow cytometry to detect death in G) CD3+T cells and H) AT-3 cells co-cultured with TCM-activated platelets and Crizanlizumab. *, *P* < 0.05; **, *P* < 0.01; ***, *P* < 0.001

### Crizanlizumab Prevents T Cell Exhaustion Induced by Platelets from Patients with Metastatic TNBC

TAPs have been implicated in immune suppression, but their effects in human cancer remain unclear. In this experiment, we tested whether platelets derived from patients can suppress T cells and whether this suppression can be reversed by P-selectin inhibition. To determine whether the immunosuppressive role of tumor-activated platelets is conserved in human disease, we evaluated the effects of platelets isolated directly from patients with metastatic triple-negative breast cancer (mTNBC). T cells were isolated from a healthy donor, activated two days prior to the experiment, and co-cultured for 48 hours with platelets from either a different sex-matched healthy donor, mTNBC patients, or mTNBC platelets pre-treated with Crizanlizumab (**Figure 5A).** Strikingly, co-culture with mTNBC patient platelets led to a profound reduction in effector-associated marker expression in both CD4⁺ and CD8^+^ effector T cells populations compared to those T cells co-cultured with healthy donor platelets (**Figure 5B&C**). This confirms that TAPs from patients are potent suppressors of T cell immunity. Notably, treatment with Crizanlizumab during co-culture fully preserved effector functionality in both CD4⁺ and CD8⁺ subsets, restoring levels comparable to those observed with healthy platelets.

**Figure 5.**
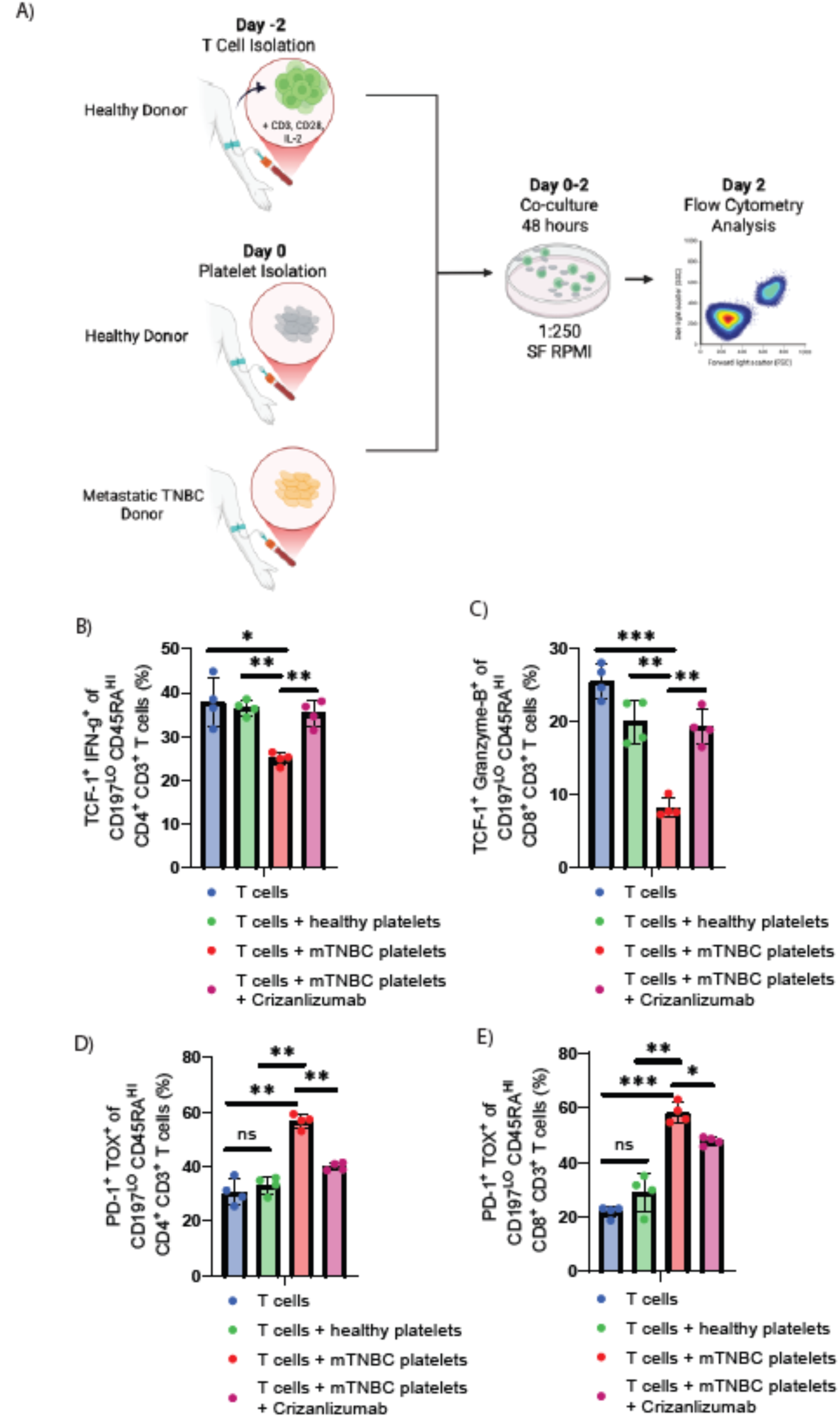
Retained functionality in human CD4+and CD8+Effector exhausted by TCM-platelets through introduction of Crizanlizumab. A) Schematic of isolation and T cells co-cultured with platelets from health and Metastatic Triple Negative Breast Cancer (TNBC) donor experiment. Flow cytometry used to quantify functionality of B) CD4+ and C) CD8+ effector T cells and exhaustion of D) CD4^+^ and E) CD8^+^effector T cells in TNBC co-culture with Crizanlizumab. *, *P* < 0.05; **, *P* < 0.01; ***, *P* < 0.001.

Moreover, mTNBC platelets robustly induced exhaustion, as evidenced by increased co-expression of PD-1 and TOX in both CD4^+^ effector T cells and CD8^+^ effector T cells (**Figure 5D–E**). However, in the presence of Crizanlizumab, this exhaustion phenotype was significantly attenuated—demonstrating that P-Selectin inhibition is sufficient to block platelet-mediated immunosuppression even when platelets are derived from patients with advanced cancer. This strongly supports the therapeutic potential of targeting the P-Selectin/PSGL-1 axis to preserve T cell functionality in mTNBC and potentially other cancers.

### P-Selectin inhibition enhances ICI response in TNBC tumors

Therefore, we evaluated RB40.34—the antibody used in the FDA approval process for crizanlizumab [65] —for its ability to block P-selectin and impact tumor growth in vivo. We first tested whether the combination of RB40.34 with anti-PD-1 would suppress tumor growth *in vivo*. Mice injected with either AT-3 or EO771 murine TNBC cells that received RB40.34 + anti-PD-1 had a significantly lower tumor volume than mice that received the monotherapies or sham (**Figure 6A**, **Supplementary Tables 4, 5)**. Flow cytometric analyses showed that mice that receive RB40.34 + anti-PD-1 had significantly higher number of functionally active CD4^+^ and CD8^+^ effector T cells found (**Figure 6B**) in the spleen. Consistently with what was found in the *in vitro* experiments with Crizanlizumab, there was significantly decreased exhaustion found in CD4^+^and CD8^+^ effector T cells in mice that received the combination or RB40.34 + anti-PD-1 (**Figure 6C and Supplementary Figure 4A-B**). We also identified pre-exhausted CD8^+^ and CD4^+^ effector T cells in the spleens of the mice that received RB40.34 + anti-PD-1 treatment (**Supplementary Figure 4C-D**) and differences in the spleens of the myeloid cell populations which has previously been reported to also express PSGL-1[66] (**Supplementary Figure 4E-F**). In addition to increase functional and decreased exhausted CD8^+^ and CD4^+^ effector T cells, we also found a significant decrease in regulatory T cells in the spleens of mice that received the combination therapy (**Figure 6D**). These results suggest that the combination therapy resulted in enhanced effector phenotypes, reduced immunosuppressive cell types, and significantly improved outcomes.

**Figure 6.**
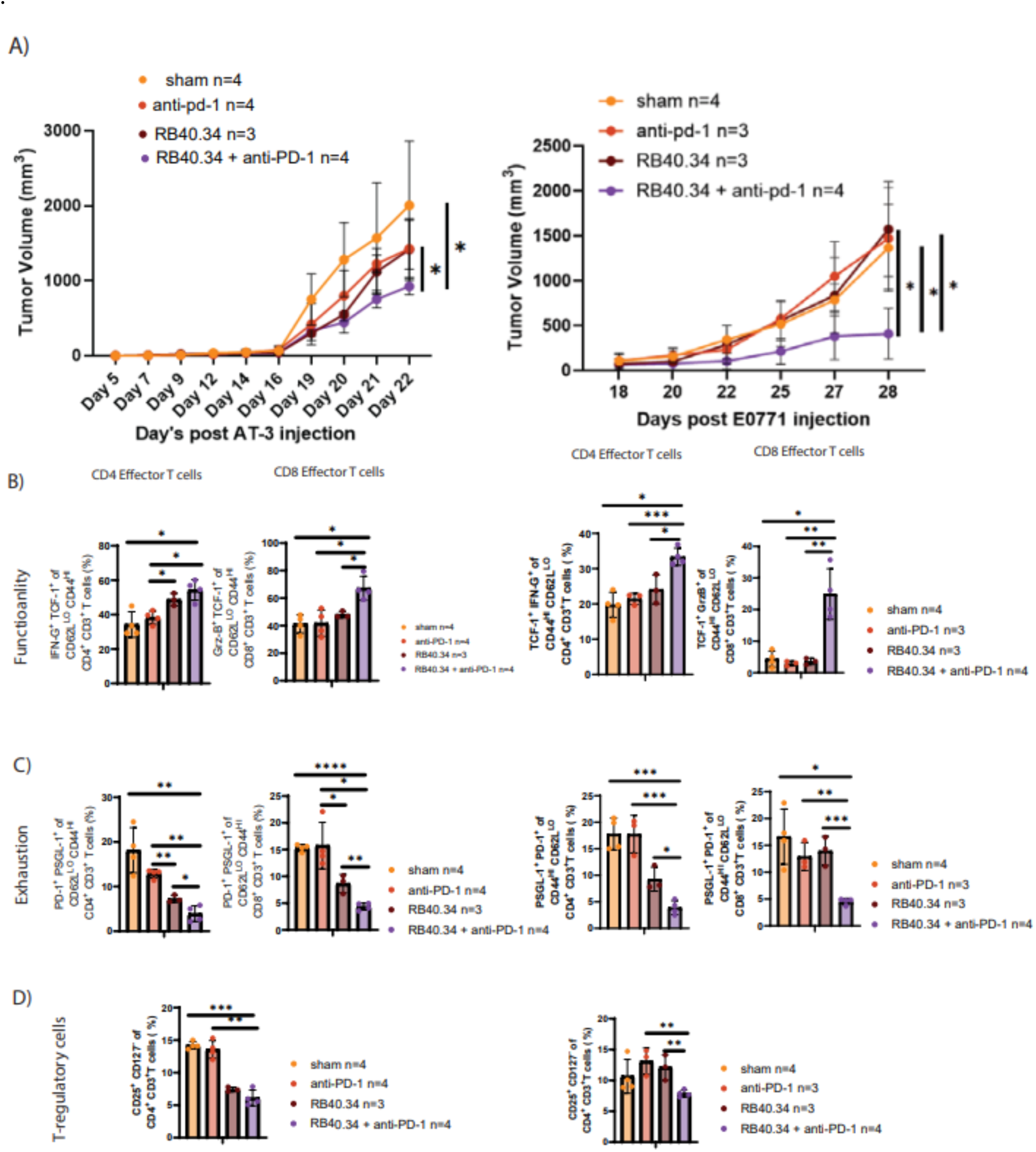
Reduced P-selectin levels increase response to ICI in tumor cells. A) Tumor volume measurements of mice injected with either AT-3or EO771 that received combination of RB40.34 and anti-PD-1. Flow cytometry measured functionally active B) CD4+, and CD8+ effector T cells co-cultured with RB40.34 and anti-PD-1, exhaustion in C) CD4+and CD8+effector T cells co-cultured with RB40.34, and D) T-regulatory cells. *, P < 0.05; **, P < 0.01; ***, P < 0.001.

## Discussion

These findings reveal a previously underappreciated role for platelets in shaping systemic anti-tumor immunity. Specifically, platelets pre-treated with conditioned media collected from AT-3 or MDA-MB-231 tumor cells derived effector T cell exhaustion and reduce cytotoxic function, thereby limiting T cells’ ability to kill tumor cells. Importantly, these effects were observed in T cells and platelets isolated from peripheral blood, demonstrating that tumor-activated platelets can induce systemic T cell dysfunction, with potential consequences for disease progression and opportunities for therapeutic intervention. By impairing circulating T cell effector function, tumor-activated platelets may promote systemic immunosuppression and tumor dissemination, suggesting that interventions targeting platelet–T cell interactions could enhance antitumor immunity and improve patient outcomes[2, 67, 68]. Circulating immune suppression may influence metastatic spread, limit disease recurrence, and undermine the efficacy of adoptive T cell therapies, all of which depend on intact systemic immunity.

Tumor cell conditioned media (TCM), which is rich in soluble factors secreted by tumor cells, appears to reprogram platelets into active modulators of immune suppression. For this study, we selected TCM from the AT-3 cell line, an ICI-non-responsive and highly aggressive model, to better understand these interactions[69]. This modulation occurs through mechanisms such as the upregulation of surface molecules like P-Selectin, which enhances platelet-leukocyte interactions, and PD-1, which can drive T cell exhaustion These findings highlight a novel mechanism by which TAPs directly interfere with T cell function, offering new insights into immune escape mechanisms in cancer.

In TNBC, platelets play critical roles in hemostasis by maintaining normal clotting, but tumor-activated platelets can become hyperactive, contributing to cancer-associated thrombosis, as well as promoting metastasis, immune evasion, and therapy resistance [24, 70–72]. Together, these data support a model in which platelet-derived P-selectin promotes T cell dysfunction via PSGL-1 engagement and suggest that disrupting this axis may be a promising strategy to restore anti-tumor immunity in TNBC.

We discovered a significantly increase amounts of functional CD4^+^ and CD8^+^ effector T cells in our in vitro and in vivo data. In vitro, Crizanlizumab helped preserve T cell function and cytotoxicity. However, in vivo the monotherapy of RB40.34 had comparable results to anti-PD-1 treatment. This data mimics more similarly the P-Selectin KO platelet data in which it appears that potentially in the absence of P-Selectin, platelets may still exhaust T cells through the PD-1/PD-L1 pathways. Crizanlizumab specifically binds to the N-terminal domain of P-Selectin while RB40.34 binds to the lectin domain of P-Selectin, which may lead to some differences [73] [74]. However, the *in vivo* data show that the blockade of P-Selectin or PD-1 plays a role in tumor growth compared to the SHAM treatment but that the combination of P-Selectin and PD-1 blockade has a greater response in an ICI-nonresponsive TNBC cell line. In the combination group alone, we identified pre-exhausted T cells were significantly upregulated compared to other treatment types. The subtypes of T cells have been shown to be a key population that responds to ICI and possess stem-like characteristics [75–79]. On top of the pre-exhausted T cells found, we also identified increase functionality and decrease exhaustion of effector T cells in the combination group that received both PD-1 and P-Selectin blockade showcasing that the combination therapy does help prevent T cell exhaustion.

While anti-platelet therapies such as aspirin have shown promise, their inconsistent results underscore the need for precise strategies that selectively disrupt platelet-tumor interactions without impairing normal platelet function. Moreover, platelet-derived factors, including cytokines and growth factors, can influence T cell activation, potentially reshaping the immune microenvironment and impacting immune responses [80]. Understanding these mechanisms could inform novel therapeutic approaches that mitigate platelet-mediated immunosuppression while preserving hemostatic balance, ultimately enhancing the efficacy of immunotherapies in TNBC.

Although platelets are well recognized for their role in driving tumor growth, our findings provide direct patient-derived evidence that they can also function as potent modulators of adaptive immunity [71]. Importantly, by using healthy donor T cells, we demonstrate that platelets isolated from mTNBC patients can directly induce T cell exhaustion *independent of tumor cells*, underscoring the novel concept that TAPs alone are sufficient to drive this dysfunctional phenotype. Notably, platelets from TNBC patients exhibit upregulation of immunomodulatory molecules, including P-selectin and PD-L1, which likely mediate this suppressive effect [81]. Beyond illuminating immune suppression mechanisms, our study addresses the critical need for predictive biomarkers of ICI response in TNBC. Existing biomarkers—such as PD-L1 expression or tumor-infiltrating lymphocyte (TIL) density—have limited sensitivity in stratifying responders [82–85]. Given their immunoregulatory role, TAP-derived proteins, platelet-expressed PD-L1, and P-selectin expression profiles may offer accessible, blood-based biomarkers for monitoring immunotherapy responsiveness [71, 86–88].

These data reveal that TAPs are not passive bystanders but active immune regulators capable of undermining anti-tumor immunity. Moreover, given that ICIs targeting PD-L1 could also alter platelet phenotype and function, platelet–T cell interactions may represent an underappreciated variable influencing therapeutic efficacy. Understanding and disrupting this platelet-driven immune suppression could open new avenues for restoring robust T cell responses while maintaining normal platelet function in cancer patients.

Despite these insights, several limitations warrant consideration. Although our in vitro TCM models do not replicate the full complexity of the tumor microenvironment, where platelet activation is shaped by multiple systemic and local factors, these experiments show that tumor-derived signals alone can activate platelets and suppress effector T cells in the circulation. Nevertheless, our *in vitro* data, demonstrated that the blockade of P-selection in combination of anti-PD-1 significantly reduced the tumor burden in different preclinical TNBC models **(Figure 6A)**. While our findings indicate that P-Selectin inhibition can restore T cell function in TNBC, the efficacy of this strategy in other cancer types remains unknown. Additional studies are required to determine whether targeting platelet–T cell interactions could serve as a broadly effective immunotherapeutic strategy across multiple cancer types. Further investigation is also needed to dissect how platelets and T cells interact within the human tumor microenvironment, including the roles of cytokines, integrins, and growth factors. Elucidating these pathways may enable strategies that selectively disrupt platelet-driven immunosuppression while preserving hemostatic balance. Together, these findings position platelets as active regulators of anti-tumor immunity in TNBC and as promising targets for enhancing immunotherapy efficacy.

