## Supplementary figures and images for "Platelet-Mediated Suppression of T Cell Function Drives Immune Evasion in Triple Negative Breast Cancer through the P-Selectin / P-Selectin glycoprotein ligand-1 Pathway"

### Supplementary Figure 1

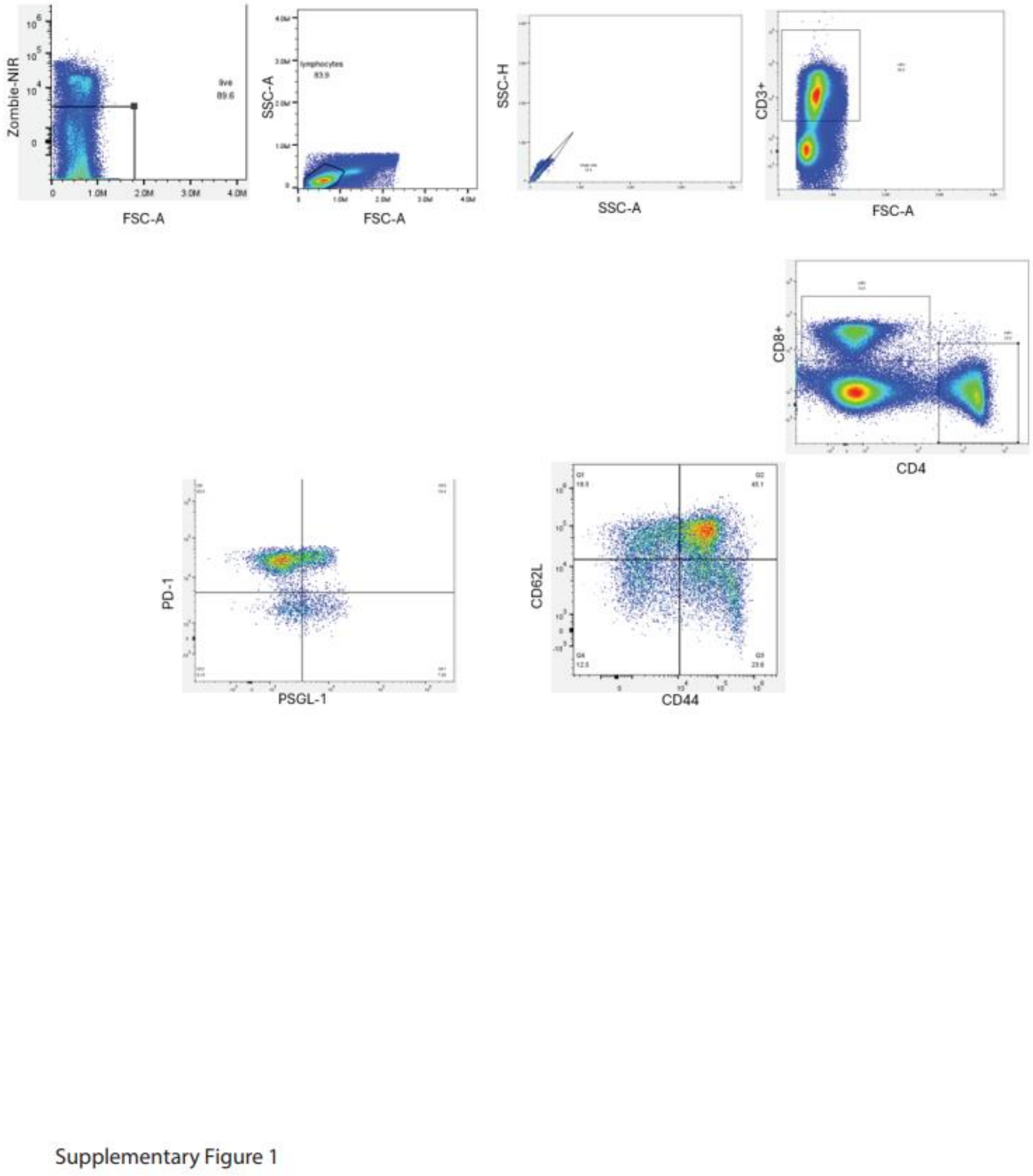


Supplementary Figure 1:

Gating strategies used to identify different T cell types in mouse.

### Supplementary Figure 2

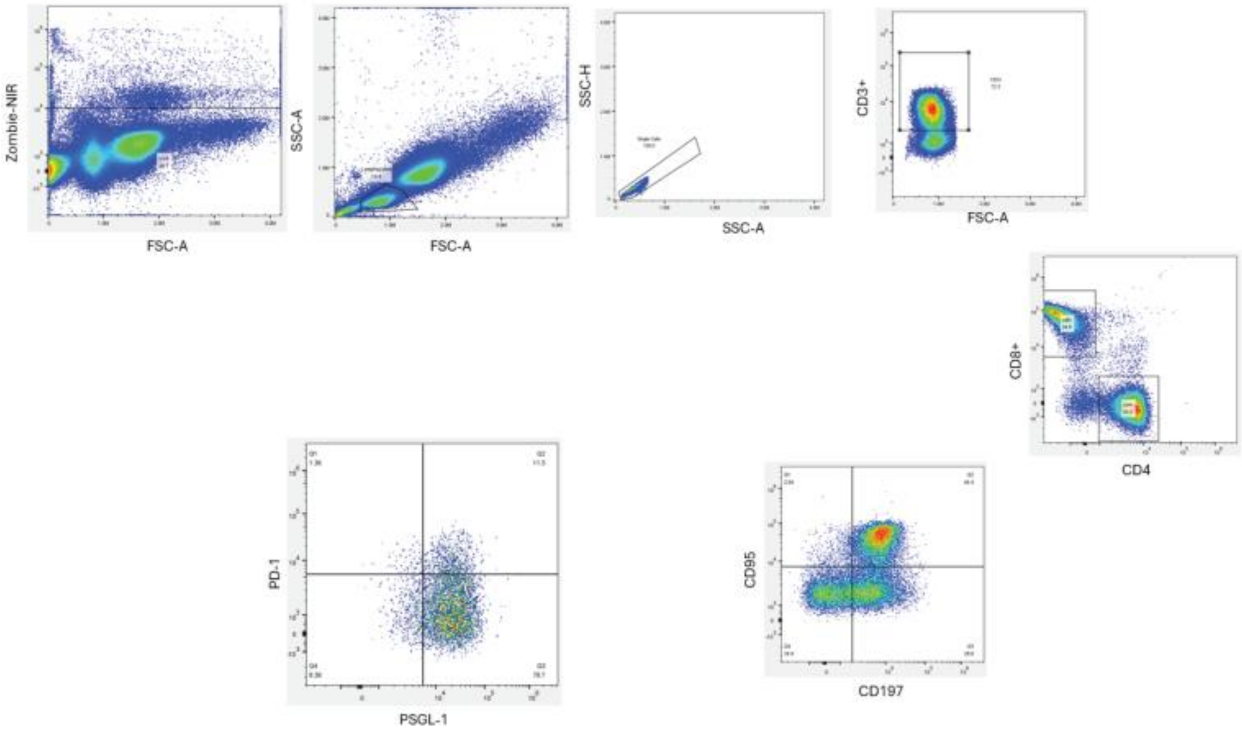


Supplementary Figure 2:

Gating strategies used to identify different T cell types in human.
