## Supplementary Figure 3 for "Platelet-Mediated Suppression of T Cell Function Drives Immune Evasion in Triple Negative Breast Cancer through the P-Selectin / P-Selectin glycoprotein ligand-1 Pathway"

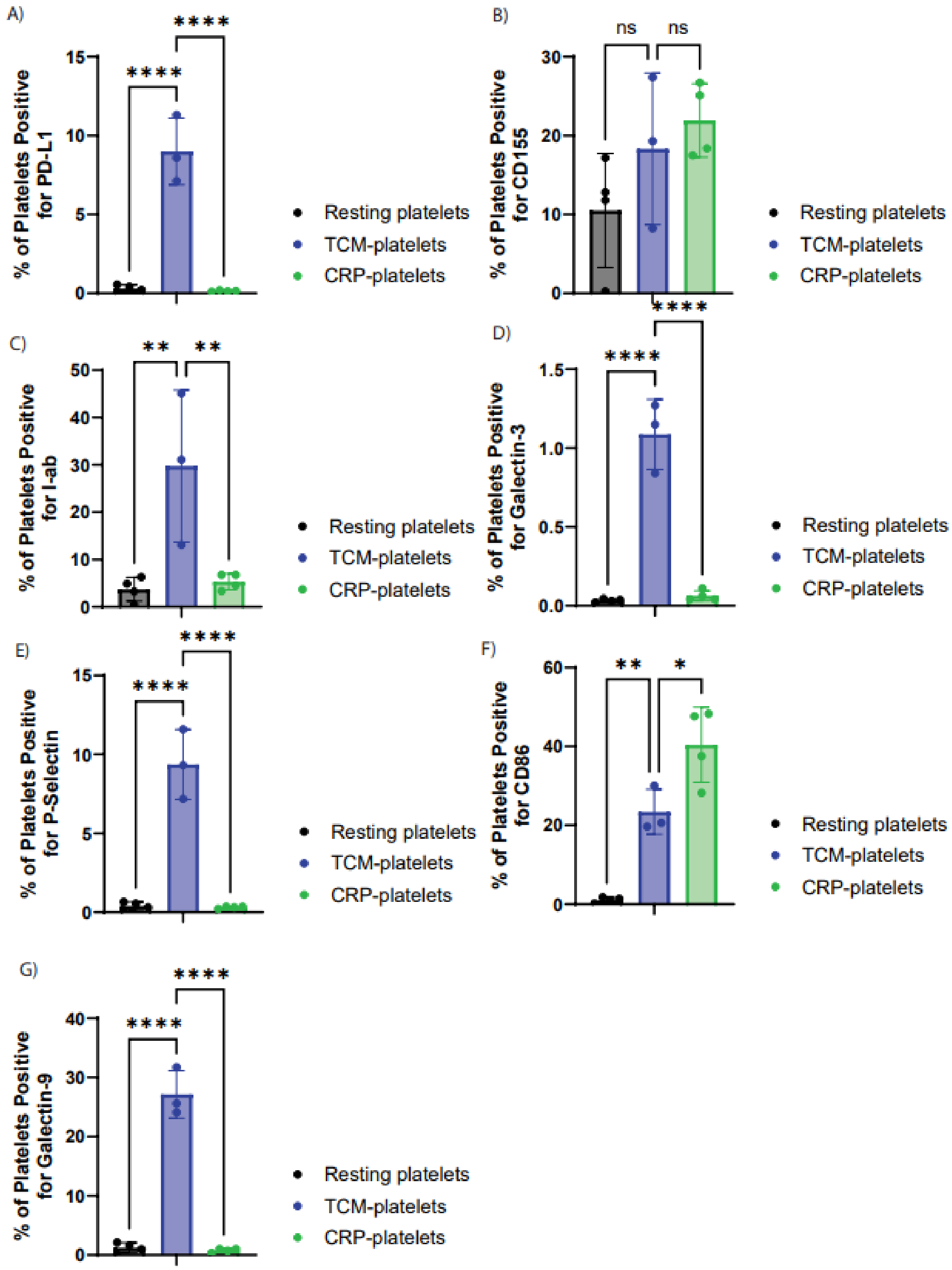


Supplementary Figure 3: Platelet expression of immune regulatory molecules. Expression from resting, TAPs, collagen related peptide (CRP)-activated platelets of A) PD-L1 B) CD155 C) P-Selectin D) CD86 E) Galectin-3 F) Galectin-9 G) I-ab. *, $P < 0.05$; **, $P < 0.01$; ***, $P < 0.001$.
