## Supplementary Figure 4 for "Platelet-Mediated Suppression of T Cell Function Drives Immune Evasion in Triple Negative Breast Cancer through the P-Selectin / P-Selectin glycoprotein ligand-1 Pathway"

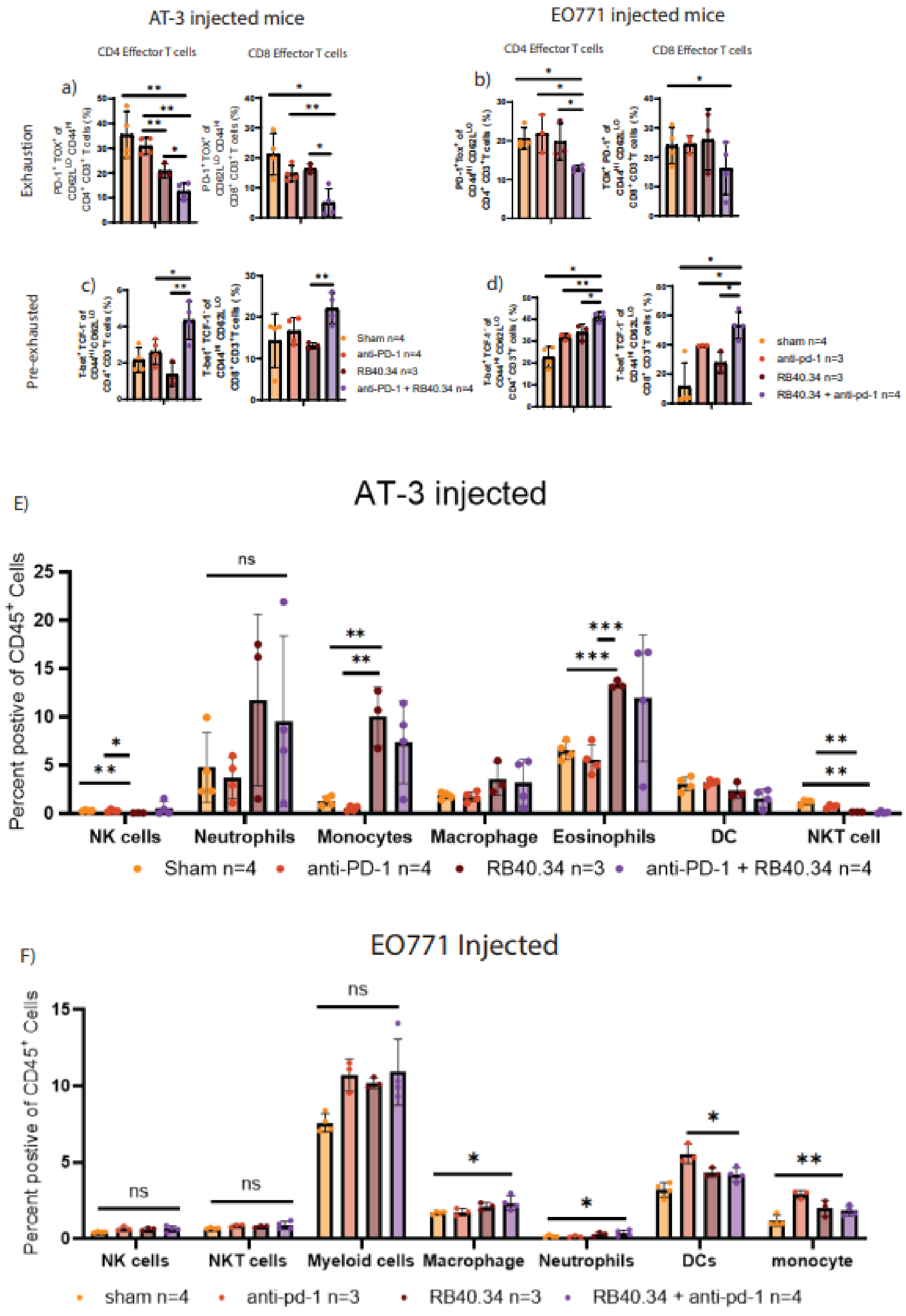


Supplementary Figure 4: Title. Flow cytometry performed to measure exhaustion in CD4 A) and CD8 B) effector T cells and pre-exhaustion in CD4 C) and CD8 D) in AT-3 and E0771 injected mice. Other myeloid cell populations of E) AT-3 injected mice and F) E0771 injected mice were measured with flow cytometry. *, $P < 0.05$; **, $P < 0.01$; ***, $P < 0.001$.
