## Supplemental Table 1 for "Platelet-Mediated Suppression of T Cell Function Drives Immune Evasion in Triple Negative Breast Cancer through the P-Selectin / P-Selectin glycoprotein ligand-1 Pathway"

| Name | Conjugate | Brand | Catalog Number | Clone | Dilution Factor |
|---|---|---|---|---|---|
| CD107a | PerCP-eFluor 710 | Thermo Fish | 46-1071-82 | 1D4B | 1/100 |
| CD127 | SuperBright 645 | Thermo Fish | 64-1271-82 | A7R34 | 1/100 |
| CD25 | BV785 | Biolegend | 102051 | PC61 | 1/100 |
| CD3 | APC-Cy7 | Biolegend | 100330 | 145-2C11 | 1/200 |
| CD4 | PE-Cy5 | Biolegend | 100514 | RM4-5 | 1/400 |
| CD44 | BV605 | Biolegend | 103047 | 1M7 | 1/100 |
| CD45 | Alexa Fluor 700 | Biolegend | 103127 | 30-F11 | 1/200 |
| CD62L | PE-Cy7 | Biolegend | 104418 | MEL-14 | 1/100 |
| CD8 | Spark Blue 550 | Biolegend | 100780 | 53-6.7 | 1/200 |
| Granzyme-B | Pe-Cyanine 5.5 | Thermo Fish | 35-8898-82 | NGZB | *1/100* |
| *IFNg* | *PE-Dazzle 594* | *Biolegend* | *505846* | *XMG1.2* | *1/100* |
| LAG3 | BV421 | Biolegend | 125221 | C9B7W | 1/100 |
| MHCI | FITC | Biolegend | 115104 | KH114 | 1/400 |
| PD1 | eFluor 450 | Biolegend | 48-9981-82 | RMP1-30 | 1/100 |
| PSGL-1 | BV510 | BD Biosciences | 563448 | 2PH1 | 1/1000 |
| Tbet | eFluor 660 | Thermo Fish | 50-5825-82 | eBio4B10 (4B10) | 1/20. |
| TCF1 | PE | BD Biosciences | 564217 | S33-966 | 1/20. |
| TIM3 | BV711 | Biolegend | 119727 | RMT3-23 | 1/100 |
| TOX | APC | Miltenyi | 130-118-335 | REA473 | 1/25. |
| Viability | Zombie NIR Fixable | Biolegend | 423105 | - | 1/1000 |

**Supplementary Table 1.** Mouse T-cell exhaustion flow panel
