## Supplemental Table 2 for "Platelet-Mediated Suppression of T Cell Function Drives Immune Evasion in Triple Negative Breast Cancer through the P-Selectin / P-Selectin glycoprotein ligand-1 Pathway"

| Name | Conjugate | Brand | Catalog Number | Clone | Dilution Factor |
| --- | --- | --- | --- | --- | --- |
| CD45 | BV605 | Biolegend | 103139 | 30-F11 | 1000 |
| IA/IE (MHCII) | BV650 | Biolegend | 107641 | M5/114.15.2 | 1000 |
| CD11b | APC/Cy7 | Biolegend | 101225 | M1/70 | 400 |
| CD3e | PE | Biolegend | 100307 | 145-2C11 | 400 |
| CD274-PDL1 | PE-Dazzle 594 | BD Biosciences | 568085 | 10F.9G2 | 400 |
| Ly6G | AF700 | Biolegend | 127622 | 1A8 | 200 |
| F4/80 | PE/Cy7 | Biolegend | 123113 | BM8 | 200 |
| Ly6C | PerCP/Cy5.5 | Biolegend | 128011 | HK1.4 | 200 |
| CD4 | AF488 | Biolegend | 100529 | RM4-5 | 100 |
| NK 1.1 | APC | Biolegend | 103212 | RA3-6B2 | 100 |
| CD8 | PercP | Biolegend | 100731 | 53-6.7 | 100 |
| CD64 | BV421 | Biolegend | 139309 | X54-5/7.1 | 50 |
| PSGL-1 | BV510 | BD Biosciences | 563448 | 2PH1 | 1000 |
| CD11c | BV785 | Biolegend | 117335 | N418 | 50 |
| Viability | Zombie NIR Fixable | Biolegend | 423105 | • | 1000 |

**Supplementary Table 2.** Mouse broad immune flow panel
