## Supplemental Table 3 for "Platelet-Mediated Suppression of T Cell Function Drives Immune Evasion in Triple Negative Breast Cancer through the P-Selectin / P-Selectin glycoprotein ligand-1 Pathway"

| Name | Conjugate | Brand | Catalog Number | Clone | Dilution Factor |
|---|---|---|---|---|---|
| CD8 | APC-H7 | BD | 560179 | SK1 | 0.005 |
| PSGL-1 | Super Bright 702 | ThermoFisher | CL488-65090100T | 3g8 | 0.005 |
| CD4 | APC-R700 | BD | 564975 | RPA-T4 | 0.005 |
| CD223/LAG-3 | PE/Fire™ 810 | BD | 565716 | T47-530 | 0.005 |
| CD3 | Alexa Fluor 532 | ThermoFisher | 58-0038-42 | UCHT1 | 0.005 |
| CD95 (FAS) | BV480 | BD | 746675 | DX2 | 0.01 |
| CD279 (PD-1) | PE-Cy7 | BD | 561272 | EH12.1 | 0.01 |
| TIGIT (IC) | BV421 | Biolegend | 372709 | A15153G | 0.01 |
| CD366 (TIM3) | BV786 | BD | 742857 | 7D3 | 0.01 |
| CD197 (memory/effector) | PE-Cy5.5 | ThermoFisher | 35-1979-42 | 3D12 | 0.01 |
| CD45RA (memory/effector) | BV650 | BD | 563963 | HI100 | 0.01 |
| CD25 (T-reg) | Alexa Fluor 488 | BioLegend | 356132 | M-A251 | 0.01 |
| CD152 (CTLA-4) | PE-eFluor 610 | ThermoFisher | 61-1529-42 | 14D3 | 0.02 |
| GranzymeB | eFluor450 | ThermoFisher | 48-8896-42 | N4TL33 | 0.005 |
| TCF1 | PE | BioLegend | 655208 | 7F11A10 | 0.01 |
| IFN-g | eFluor506 | ThermoFisher | 69-7319-42 | 4S.B3 | 0.01 |
| Foxp3 | Spark NIR 685 | BioLegend | 320130 | 206D | 0.01 |
| TOX | APC | Miltenyi Biotec | 130-118-335 | REA473 | 0.04 |
| Viability | Zombie-NIR | Biolegend | 423105 | • | 1/1000 |

**Supplementary Table 3.** Human T-cell exhaustion flow panel
