## Supplemental Table 4 for "Platelet-Mediated Suppression of T Cell Function Drives Immune Evasion in Triple Negative Breast Cancer through the P-Selectin / P-Selectin glycoprotein ligand-1 Pathway"

| Group Number | Mouse ID | Treatment | Mass (g) | Tumor Mass (g) |
|---|---|---|---|---|
| Control | Control 1 | No Treatment | 20.53 | 0 |
| Control | Control 2 | No Treatment | 19.19 | 0 |
| Control | Control 3 | No Treatment | 18.42 | 0 |
| Control | Control 4 | No Treatment | 18.47 | 0 |
| 1 | 330 | RB40.34 | 18.06 | 0.812 |
| 1 | 331 | RB40.34 | 21.14 | 1.852 |
| 1 | 328 | RB40.34 | 22.85 | 1.153 |
| 1 | 339 | RB40.34 | 20.27 | 0.941 |
| 2 | 340 | Anti-PD-1 + RB40.34 | 18.34 | 0.788 |
| 2 | 332 | Anti-PD-1 + RB40.34 | 20.95 | 1.02 |
| 2 | 337 | Anti-PD-1 + RB40.34 | 20.34 | 1.045 |
| 2 | 355 | Anti-PD-1 + RB40.34 | 23.01 | 0.742 |
| 3 | 341 | Anti-PD-1 | 20.08 | 1.791 |
| 3 | 335 | Anti-PD-1 | 20.7 | 1.052 |
| 3 | 333 | Anti-PD-1 | 20.85 | 1.51 |
| 3 | 329 | Anti-PD-1 | 18.37 | 0.392 |
| 4 | 334 | Sham | 19.97 | 1.371 |
| 4 | 338 | Sham | 19.8 | 1.772 |
| 4 | 336 | Sham | 19.67 | 1.41 |
| 4 | 342 | Sham | 20.12 | 1.576 |

**Supplementary Table 4.** AT-3 injected mouse tumor data
