## Supplemental Table 5 for "Platelet-Mediated Suppression of T Cell Function Drives Immune Evasion in Triple Negative Breast Cancer through the P-Selectin / P-Selectin glycoprotein ligand-1 Pathway"

| Group Number | Mouse ID | Treatment | Mass (g) | Tumor Mass (g) |
|---|---|---|---|---|
| PBS injected | Control 1 | No Treatment | 19.78 | - |
| PBS injected | Control 2 | No Treatment | 18.17 | - |
| PBS injected | Control 3 | No Treatment | 18.53 | - |
| 1 | 405 | SHAM | 21.69 | 1.4049 |
| 1 | 450 | SHAM | 21.47 | 0.776 |
| 1 | 419 | SHAM | 19.95 | 0.665 |
| 1 | 449 | SHAM | 19.59 | 0.991 |
| 2 | 443 | Anti-PD-1 | 21.62 | 1.291 |
| 2 | 444 | Anti-PD-1 | 23.2 | 1.229 |
| 2 | 419 | Anti-PD-1 | 21.11 | 0.684 |
| 3 | 446 | RB40.34 | 20.53 | 1.504 |
| 3 | 413 | RB40.34 | 19.92 | 0.858 |
| 3 | 415 | RB40.34 | 21.78 | 0.91 |
| 4 | 447 | Anti-PD-1 + RB40.34 | 22.49 | 1.27 |
| 4 | 442 | Anti-PD-1 + RB40.34 | 21.74 | 1.025 |
| 4 | 411 | Anti-PD-1 + RB40.34 | 18.09 | 0.48 |
| 4 | 410 | Anti-PD-1 + RB40.34 | 20.45 | 0.45 |

**Supplementary Table 5.** EO771 injected mouse tumor data
